# Quantitative Description of *C. elegans* mRNA Landscapes From High Coverage Single-Cell Transcriptomes

**DOI:** 10.64898/2026.07.18.734896

**Authors:** Florian Bernard, Emma Kandel, Delphine Dargere, Eric Cornes, Denis Dupuy

## Abstract

Single-cell RNA sequencing technology dramatically changed the way we investigate transcriptomes. However, the amount and complexity of data generated by such methods poses new challenges for biologists who are trying to extract detailed insights into the genetic programs that drive cellular functions and differentiation. To provide a more intuitive understanding of cell specific gene expression programs, we developed a novel approach for exploiting scRNA-seq data that detects individual gene expression levels in each cell, by avoiding dimensional reduction methods. This was achieved by focusing our analysis on individual cells with a high sequencing coverage (above 15000 Unique Molecular Identifiers (UMIs)). Such High Coverage Cells (HCC), were found in all five *C. elegans* scRNA-seq datasets we investigated and constitute direct quantitative experimental observations of the mRNA content of individual cells. Clustering the complete gene expression matrix for these cells, we identified gene sets specific for most *C. elegans* tissues. Among each set we found genes that are dominating cell specific transcriptomes as well as genes that are restricted to particular cell types but are a thousand fold less expressed. For each cell type or subtype we characterized, we identified a set of genes with expression restricted to those cells that were not previously associated with the corresponding tissue. Our results demonstrate that by focusing on HCCs, we can provide high-resolution quantitative descriptions of cellular expression landscapes that are immediately exploitable for researchers to generate new biological hypotheses. Overall, we demonstrate that HCCs represent a powerful and largely unexplored source of biological insights and suggest that future scRNA-seq experiments could benefit from focusing on HCC enrichment to capture and exploit the full complexity of cellular transcriptomes.

## INTRODUCTION

Single-cell gene expression profiling provides a rich and systematic characterization of complex systems which revolutionized the way biologists can interrogate transcriptomes^1,2^. The ability to decipher the complete transcriptional information from isolated cells offer the potential to reach detailed molecular characterization of the genetic programs that drive cellular differentiation and pathogenesis. However, the amount and complexity of data generated by such methods pose new challenges for biologists who are trying to extract biological insights on cellular functions^3-5^.

Since the first published single-cell transcriptomics study in mouse germ cells^1^ technological developments have fueled a rapid increase in the number and types of cells that can be studied in single-cell RNA-seq experiments. Large scale single-cell reference “atlases” are currently generated at a fast pace, featuring profiles of millions of cells across tissues and organs, in multiple species, during development or aging, in physiological or pathological states, and under environmental or genetic perturbations^7-9^. These reference atlases constitute a very powerful resource to shed light on fundamental questions of biology and medicine, including the origin and development of diseases, and the molecular determinant of cell types and their interactions^10^.

In their seminal 2011 publication, Islam and collaborators obtained 241,000 mRNA molecules per cell for ~90 individual cells^2^. By contrast, the currently prevalent experimental strategy involves the sequencing of libraries from tens of thousands of individual cells with a median cut-off for inclusion of 500 Unique Molecular Identifiers per cell (UMI/cell)^11,12^. To manage the complexity of the data collected, dimensional reduction methods, such as t-SNE and UMAP, project the cellular expression landscapes onto “non interpretable” space where genetic information cannot be easily accessed^3,4^. Moreover, after the initial clustering, current analysis methods stop examining cells as separate entities and instead consider gene enrichment at the level of clusters of cells. This aggregation step is made necessary because only a fraction of the transcriptome of each individual cell is actually sampled. A median of ~600 UMI/cell, while currently considered a high standard^12^, only probes less than one percent of the number of transcripts expected to be present in a cell. Here, we are exploring an alternative strategy to avoid all these complex computational steps. We reasoned that focusing on the cells with the highest-coverage (UMis/cell) would provide experimental high resolution description of cellular mRNA landscapes.

## RESULTS

### scRNAseq datasets contain an exploitable fraction of “High Coverage Cells”

Single-cell RNA sequencing (scRNA-seq) involves Illumina sequencing of cDNA libraries generated from the collection of hundreds of thousands of microfluidic droplets containing gel beads with barcoded oligonucleotides for reverse transcription priming^18^. To avoid having multiple cells associated with the same barcode, most beads are not associated with cells and only yield few mRNA reads likely coming from droplets receiving traces of cytoplasm from broken cells, often referred to as “ambient RNA background”. It is common practice to set a minimal threshold of UMI content ~500 to exclude the empty droplets, this leads to datasets that contain tens of thousands of cells, making it impossible to cluster the data in a two dimensional cell vs gene matrix.

We selected five scRNA-seq datasets from *C. elegans* (Table 1)^10-14^. We first looked at the coverage per cell in those datasets by plotting the number of cells with each observed number of Unique Molecular Identifier (UMIs) (Figure 1A). In each dataset we found a similar heavy tail with cell coverage in the hundreds of thousands of UMIs. Such a distribution could be expected since the sequencing step is sampling DNA molecules from a pool constituted from >99% of “empty” droplets. A simple generative model reproduces remarkably well the shape of UMI frequencies from individual scRNA-seq runs: we simulated variability in RNA capture by stochastic sampling to generate observed UMIs from a mixture of empty (ambient RNA) droplets and 0.4% cell-containing droplets. The different shapes observed in the experimental data are approximated using various minimal count of RNA capture from non-empty droplets (Supplementary Figure 1). This modelisation bolstered our confidence that the tail end of the experimental distributions represents a fraction of genuine cells with extremely high coverage. This constitutes a tremendous resource for in-depth analysis of cellular transcriptomic landscapes that has been so far overlooked by the community. In the combined datasets, we identified 4,543 High Coverage Cells (HCCs) with a coverage over 15,000 UMIs (30-fold higher than the current minimum standard for scRNA-seq) and the maximum observed coverage reached ~600,000 UMIs.

**Table 1.**
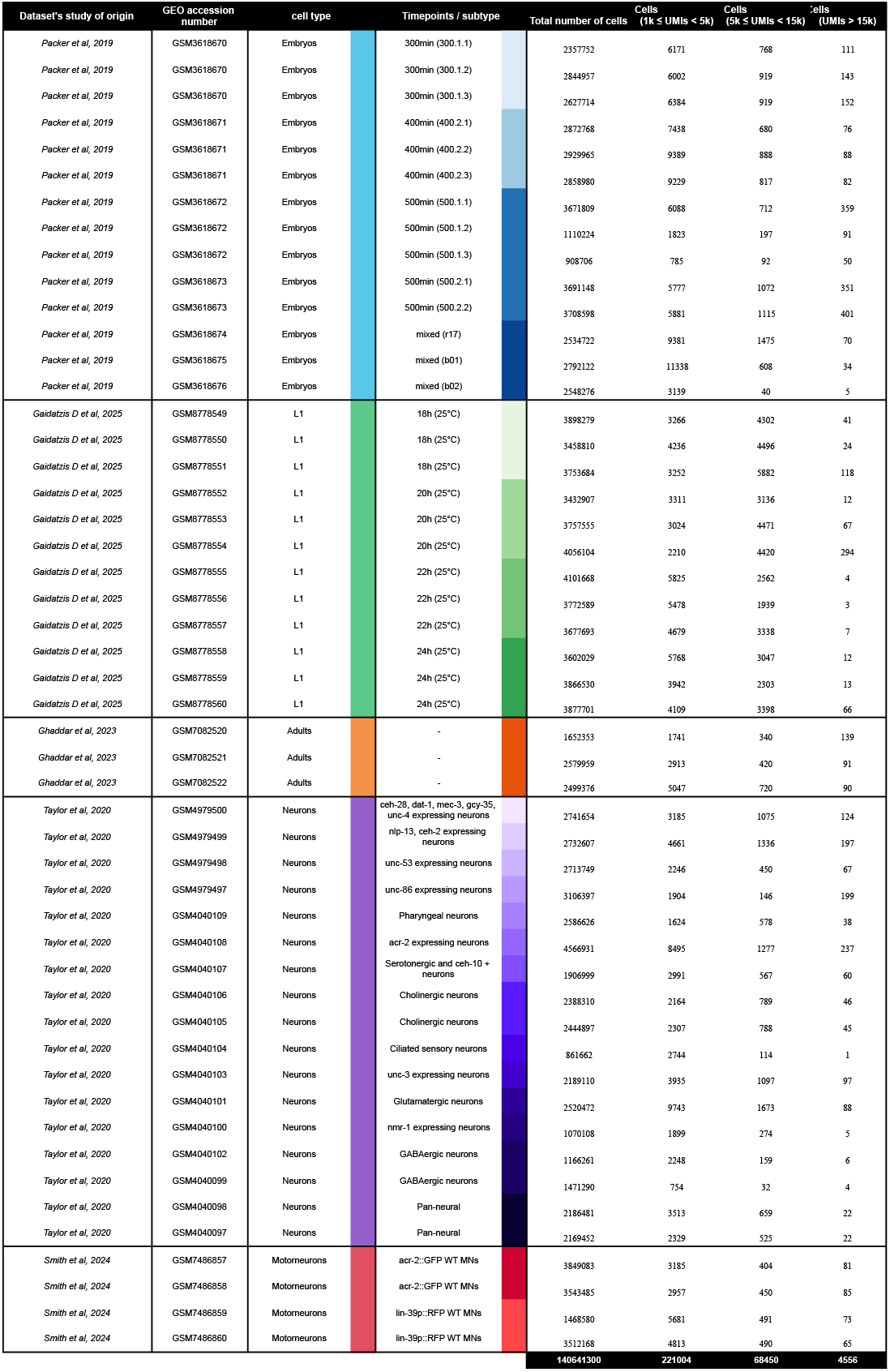
*C. elegans* scRNAseq datasets included in this analysis.

**Figure 1:**
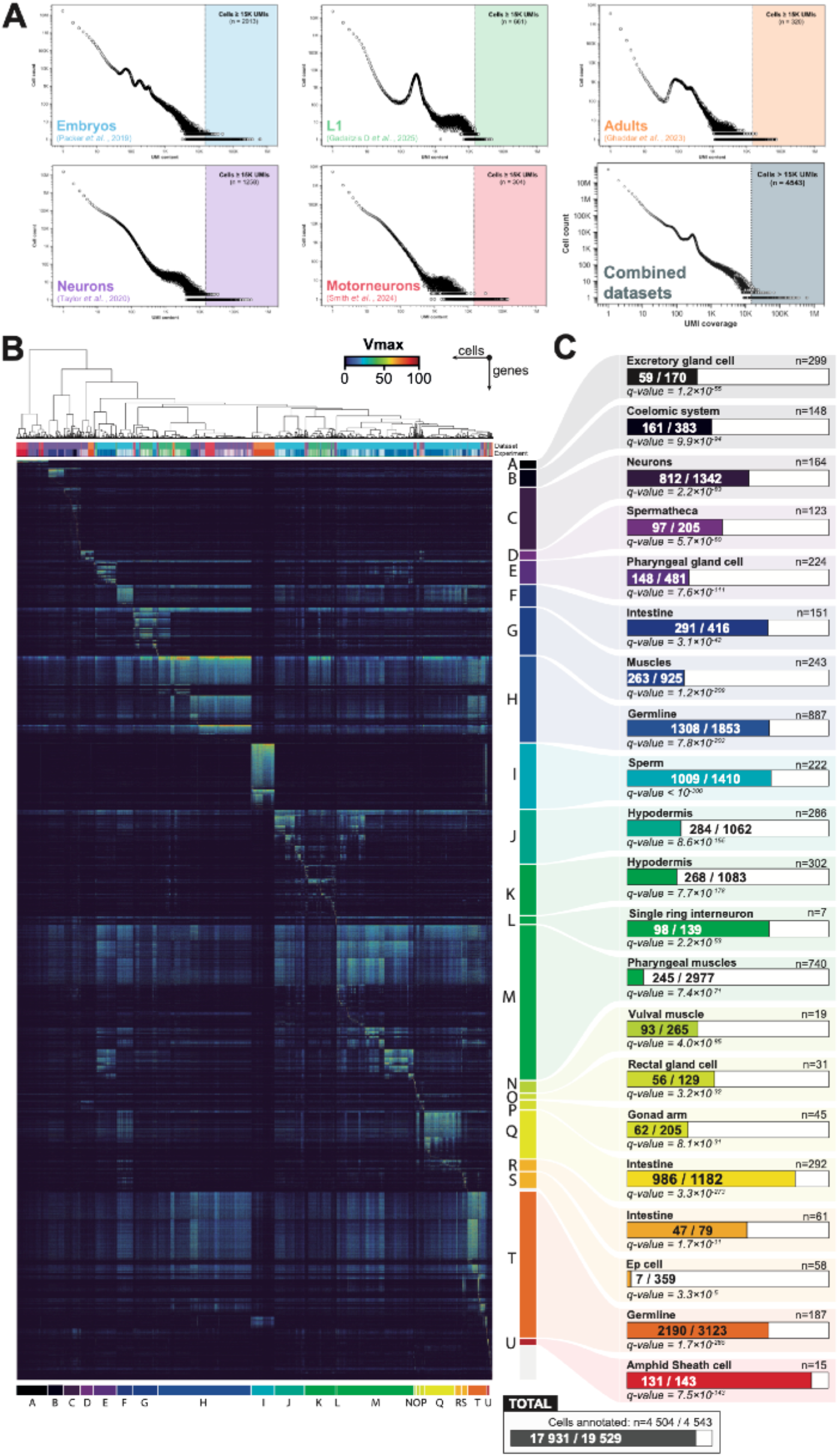
**A)** Distribution of sequencing coverage in the datasets used in this study. The shaded area indicates the high content cells selected for our analysis. **B)** Supervised clustering (see Methods) of the normalized gene expression matrix restricted to High Coverage Cells reveals distinct groups of cells characterized by clear gene clusters. Color bars indicate the dataset and individual experiment of origin of each cel according to the colors in Table 1. **C)** Tissue Enrichment Analysis of the gene clusters^15^. n= number of cells in the cluster ; fraction indicates the number of genes in the cluster associated with retained the annotation / total number of genes.

### A supervised gene clustering strategy identifies cell-type specific gene sets from HCCs

We first clustered the HCC using all 19,529 genes for which transcripts were detected in those cells, then clustered the genes according to what cell group they most correlated with. In short, we assigned cells to each of the nodes they belong to on the neighbor-joining dendrogram (see Methods). Then for each node on the tree we generated a virtual “expression profile” for a theoretical gene that would be specific to this node: with maximum expression in the cells of the node and no expression in cells outside of the node. We then compared each actual gene expression profile in our matrix with all the theoretical gene patterns using Pearson’s Correlation Coefficient as a metric, and assigned each gene to the node they matched best. We obtained a highly structured expression matrix in which gene sets specific to individual cell groups are easily identifiable (Figure 1B). This strategy gave better results than performing neighbor-joining clustering between all the genes, presumably because it is not influenced by co variation in the experimental noise. We performed tissue enrichment analysis on each of the gene lists associated with a node of the neighbor joining tree of cells, then we regrouped sibling nodes that shared a similar tissue annotation (see details in the Methods). This led us to define 21 major gene clusters that cover most of the animal’s tissues (Figure 1C). While the results of tissue enrichment analysis have an extremely robust statistical significance, it is important to bear in mind that much of the existing annotations originate from previous enrichment analyses in transcriptomic studies that include the datasets we are using here. However, the majority of genes carrying the most enriched annotation also carry additional annotations that may indicate a less restrictive expression pattern than our clustering suggests. Additionally, a significant fraction of the genes within each clusters have never been associated with the dominant tissue type so far.

### In vivo validation of tissue specificity

We selected genes from three distinct tissues with various levels of expression (Figure 2A) and performed smiFISH on *C. elegans* embryos (Figure2B). For each cluster, we selected two genes that were previously associated with the enriched tissue annotation and one that was not. We first focused on cluster F that was annotated as “Intestinal”. The most expressed gene in this cluster is *ttr-50* with a mean detection level ~100,000 rpm in the corresponding set of cells. No direct experimental investigation of this gene’s expression was available on Wormbase at this time, the various tissue annotations associated with this gene come from enrichment of its transcript in genome-wide studies and include, muscle, oocyte, mesoderm and intestine. smiFISH of *ttr-50* shows a strong signal in the embryonic intestinal cells in embryos spanning from gastrulation until the 1.5-fold stage, thus confirming the specific tissue annotation suggested by our analysis. The other two genes selected from this cluster were *cpz-1* and B0495.7 which have expression levels in the same cells averaging 10,000 rpm and 300 rpm, respectively. Both presented FISH signal in the intestinal cells at the same stages of development as *ttr-50*. For B0495.7, this is the first experimental evidence of expression in the intestinal cells. We then selected three genes (*asp-1, nep-17* and *clec-17*) from cluster Q which is also annotated as “Intestinal”, but which express a distinct set of specific genes. For these three genes, we observed a strong intestinal signal that starts at the 1.5-fold stage and persists until the 3-fold stage. We similarly selected three genes from the J and K “hypodermal” clusters and confirmed that they were indeed expressed with corresponding embryonic patterns including some temporal specificity that explains some of the differences between genes visible in some cells. Finally, we selected three genes from the “Pharyngeal Gland” cluster (E). Here again we found identical cell expression patterns in the expected cells despite the different levels of gene expression. For the limited sample we tested, in vivo it seems that the most enriched annotation emerging from our gene lists is indeed the correct one for the expression pattern. Additionally, in each tested case it seems that this annotation should be the only one retained, as no expression was detected in other tissues that had previously also been associated to those genes. Critically, this is also true for the genes that were not previously associated with their respective enriched annotation.

**Figure 2:**
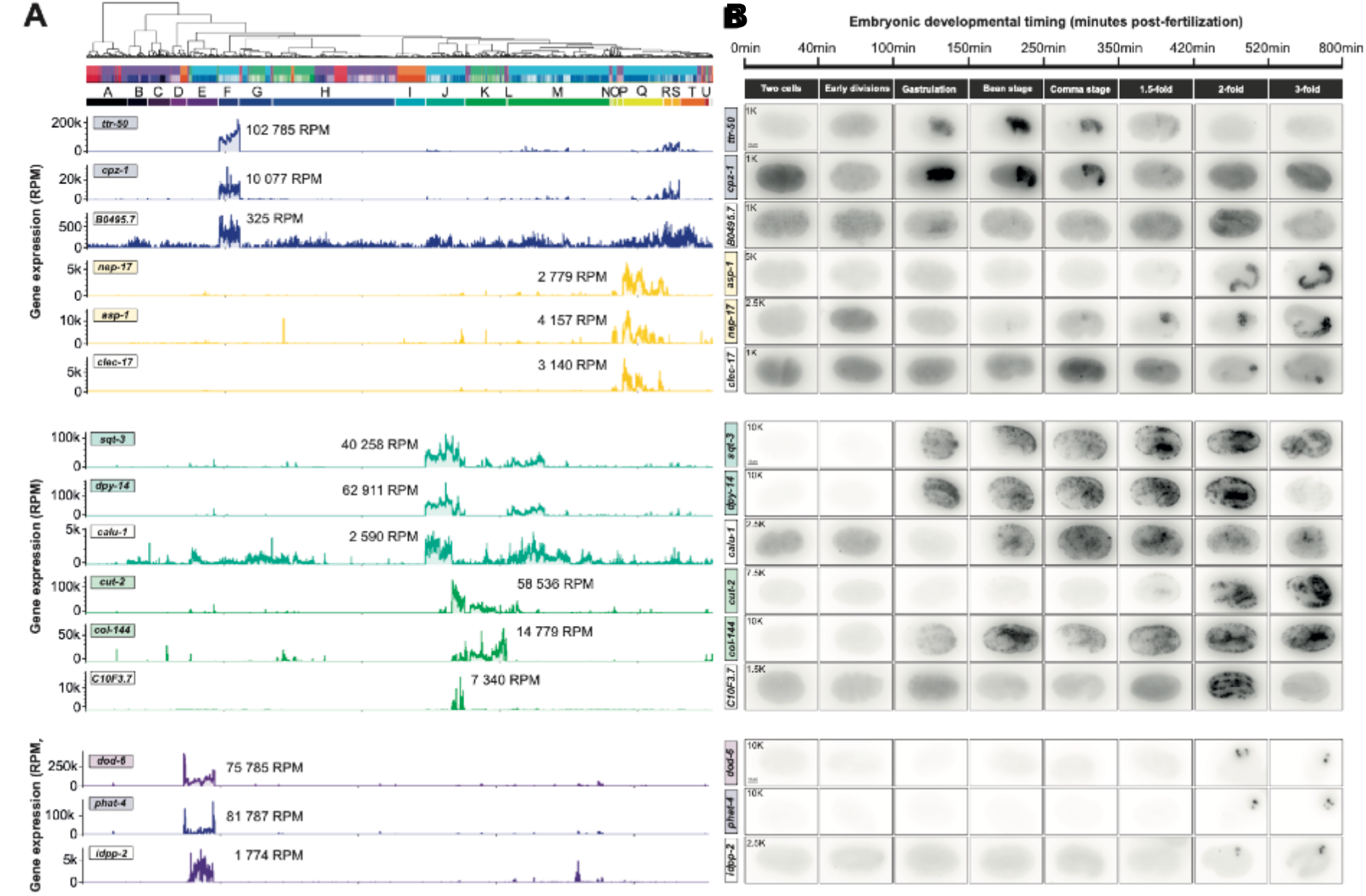
In vivo validation of tissue specificity. **A)** Expression level of individual genes across the high coverage cells ordered as in Figure 1A, the median level of expression for each gene in the cluster in indicated in rpm (reads per million). Gene names with a colored background were associated with the dominant tissue type in their respective cluster; gene names with a white background had no previous association with that tissue. B) In situ hybridization of individual genes during *C. elegans* embryonic development. For each gene we indicate the maximum value of the pixels used in the presented images.

### Cellular Gene Expression Spectrums

Most identified gene clusters contain at least one dominant gene with expression level orders of magnitude above the others (Figure 2B). We, therefore, assigned that “flared” gene name to the whole cluster as a shorthand. Based on this observation we devised a new representation for individual cells’ expression profiles, in which the relative expression level of all gene clusters can be visualized. Each gene is assigned a color according to the gene cluster they were assigned to by our supervised clustering strategy. Figure 3A presents the individual contribution (in percent) of each gene cluster to the total number of reads detected in each cell. By stacking these individual contributions we obtained a “gene clusters expression spectrum” representation that highlights the variety of gene expression programs among the High Coverage Cells (Figure 3B). Strikingly, in several annotated cells (such as excretory gland, or sperm) over 80% of the sequenced transcripts come from a small set of tissue specific genes. As we already noted, we find a large dynamic range of expression levels within each gene cluster : from a few hundred reads per million to a thousand times as many. It is therefore frequent that a single gene accounts for more than 1% of the transcripts found in a cell. Most of these “flared” genes have a strong cell type specificity and therefore constitute the most efficient cell markers when attempting to categorize cells based on their expression profiles. Thus, we generated a “Flared genes spectrum” to highlight different sub clusters of gene expression patterns within the original clusters (Figure 3C). In this representation, each gene that contributes more than 1% of the transcripts in a given cell is attributed a color, then it is plotted in the order of the clustered matrix in Figure 1B (see supplementary video). In this manner, genes with similar colors can still be distinguished by their position relative to other genes. These gene expression spectrums are a good way to visualize the dominant genes that characterize each cell type even for more specific levels of annotation found within the original clusters. Since we assigned each gene to the cell group that it matches best, some of those assignments are not always optimal. For example, cluster L is specific to a group of seven cells (Figure 1C). Among the 139 genes indicated for this cluster in figure 1C, only 83 were matched with the whole group of 7 cells (73 of these 83 genes were previously associated with the Single Ring Interneuron (RIS)). Two additional gene clusters were assigned to two subgroups of the seven cells, and also enriched for RIS (9/13 and 8/24 respectively). The expression spectrum for these cells presented in Figure 4 indicates that 3 of the cells of the group express 4 genes associated with heat shock response (F44E5.4, F44E5.5, HSP-16.48 and HSP-16.49), that are not present in the other four cells. It is possible that these differences are due to variations in experimental conditions rather than distinct developmental states of that neuron. In figure 4, we show a few additional examples of such spectrums for small sets of closely related cells. We only present a sample of the most expressed genes in the legend, in each case, that are specific to the group of cells considered. While this representation allows direct evaluation, at the macroscopic level, of the similarity between cells, it doesn’t provide the full picture. For the ASE neurons, for instance, none of the genes specific to those cells passed the threshold of 1% of the total read count required to appear on the spectrum.

**Figure 3:**
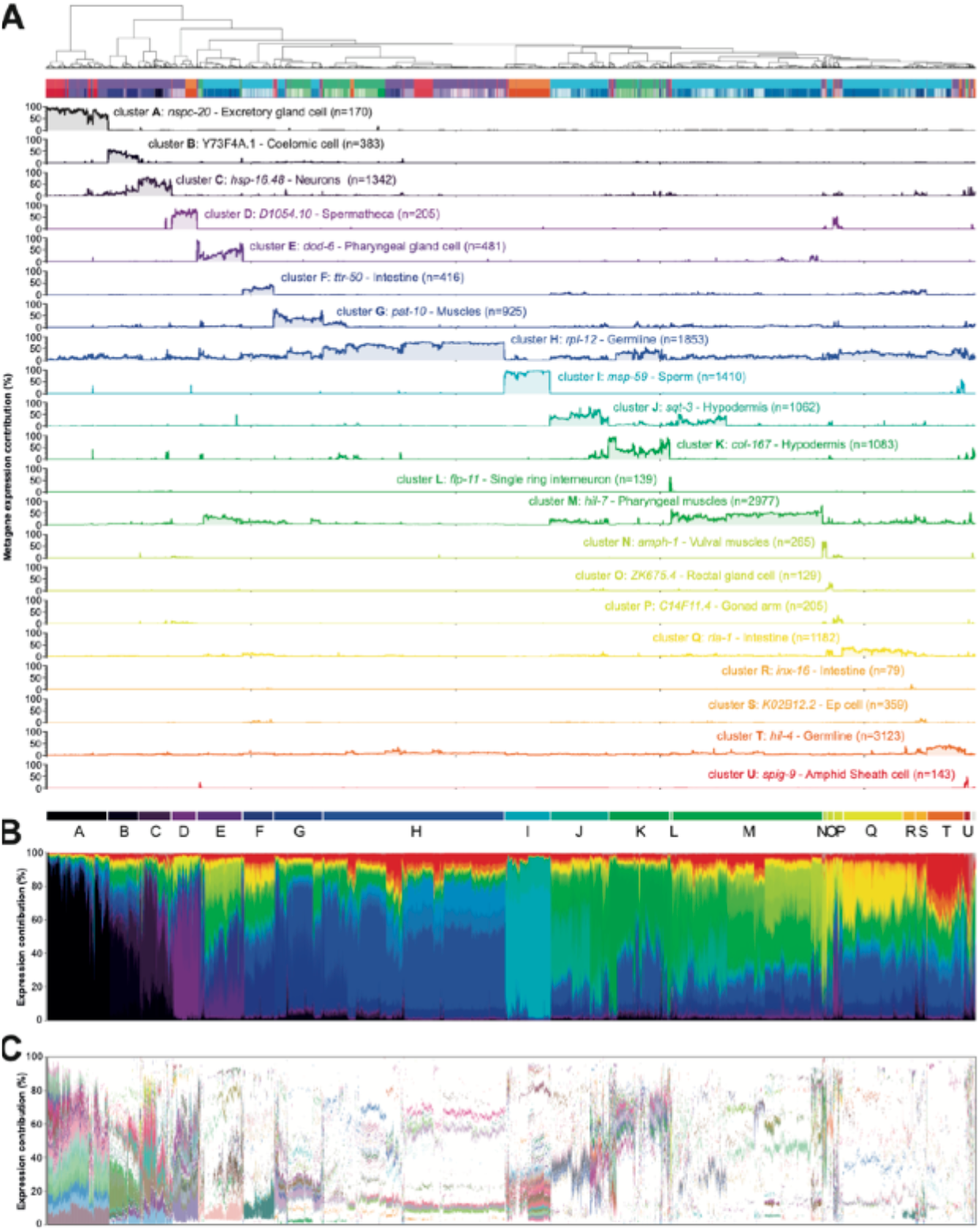
**A)** Cumulated read counts from genes in each cluster relative to the total read read counts in each cells. **B)** A “Gene clusters expression spectrum” shows the stacked gene clusters in each cell type relative to each other. **C)** Flared gene expression spectrum indicating only genes that contribute for at least 1% of the total reads in their cell (each gene is randomly assigned a distinct color, see complete legend in Supplementary Figure 2).

**Figure 4.**
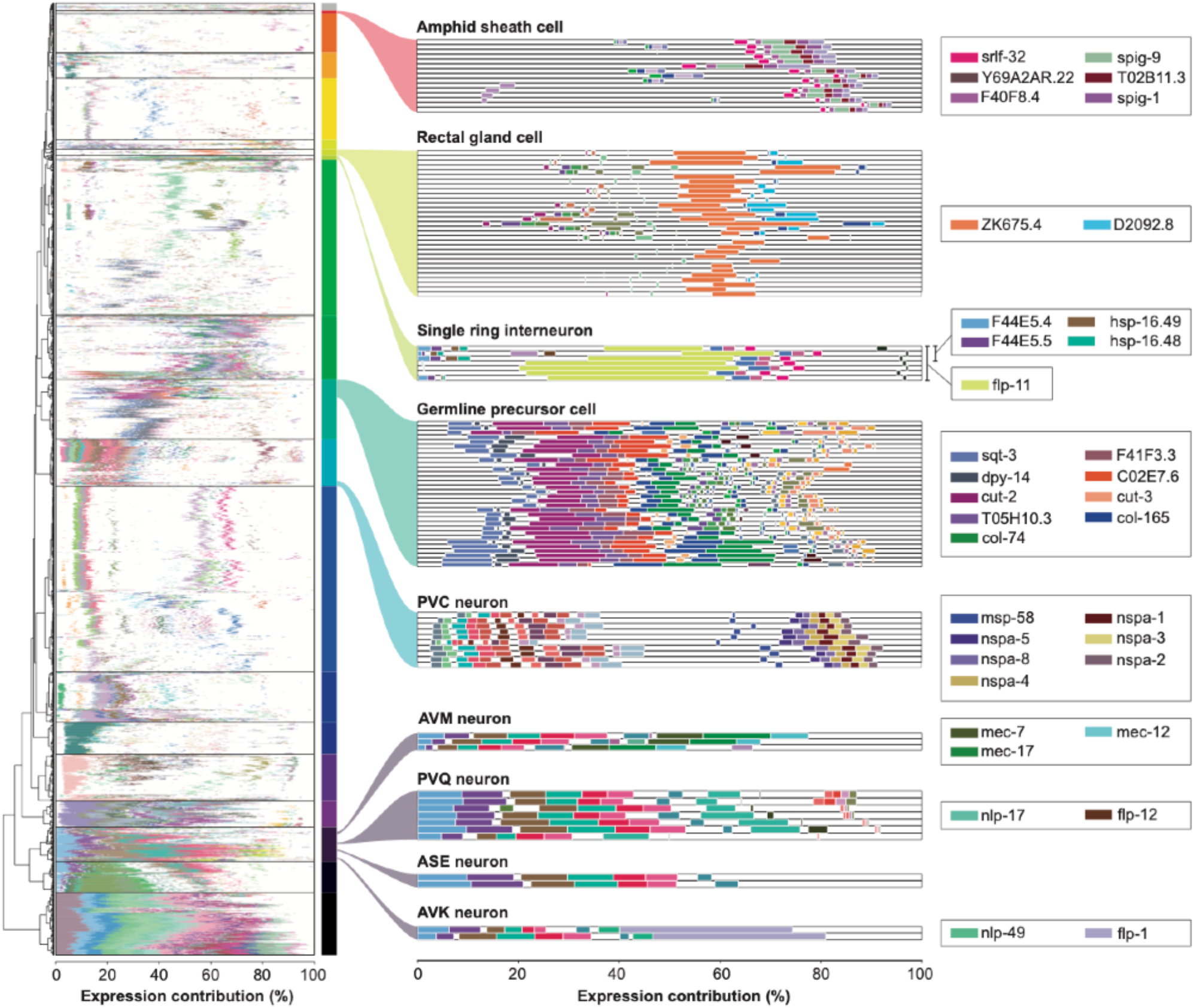
Gene expression spectrums for individual annotated cell types. The legend indicates the most discriminating flared genes for each cell type or cell group.

To validate the specificity of those genes and annotations, we can use simple expression plots. In Figure 5, we present the expression levels of the genes associated with 4 neuronal subtypes from cluster C, across the 164 cells from that cluster. The three cells annotated as “AVM” cells are dominated by three genes of the mechanosensation pathway (*mec-17, mec-7* and *mec-12*) each with a mean level of expression within the cluster between 60,000 and 107,000 reads per million, thus accounting for 25% of the reads in these cells. This strongly indicates that these cells are indeed touch receptor neurons (TRNs), however, the automatic annotation selected only AVM, ignoring the other neurons from that class that also express those genes (ALMs, PVM and PLMs). Other genes from the pathway display the same specificity, but with a hundred fold lower expression levels (*mec-1, mec-3, mec-9*), reinforcing the value of using HCC to provide quantitative measures of the relative expression of transcripts. Similar ratios are found among the genes specific for other cell types. The highest expressed gene in the “ASE” specific gene list is *C36B7*.*3*, accounting for just under 1% of the reads in those cells (Figure 5). *C36B7*.*3* has not been directly investigated to date, but has been found enriched in multiple neurons in various genome-wide studies according to Wormbase. In our HCC dataset, it is exclusively found in cells that also express *gcy-19*, a gene whose promoter has been shown to drive GFP expression only in ASE and IL2 neurons^19^. In the same gene cluster, *dsc-1* is reported to be expressed in several head neurons, including ASE, using extrachromosomal array transgenes^20^. Notably, other known markers from the ASE neurons did not associate with these cells^21^, therefore additional investigations could be carried out to validate if *C36B7*.*3* is indeed a strong specific marker for ASE. The “PVQ” annotation was derived from only 6 genes associated with these cells, however these include *nlp-17*, which was shown to be a specific marker for these cells^22^. The three genes that were not previously annotated to this neuron appear at very low intensity in only one of the seven cells, and this is the only HCC in wh i ch they were detected. Whenever a gene is found in only one cell, it will mechanically be assigned as “specific” for the group to which that cell belongs, but that association should be taken with caution. As we collect more HCC with enhanced coverage, such cases will become ra r e r and the confidence in our annotations will grow, for both cell identity and gene specificity. Finally, the case of AVK seems unambiguous as all 11 genes associated with these cells were previously found highly enriched in mRNA from sorted AVK neurons and the cellular transcripts are dominated by *flp-1*, the neuropeptide specific of that neuron^22^. Notably, all cells in the “Neuronal” cluster C show the four heat shock response genes mentioned above for a few of the RIS cells, which may reflect an experimental issue in the datasets from which they are originating from. Additional reads, such as *unc-54*, that contribute to a large fraction of the transcripts of the cell, may arise from the transgenes used in the cell sorting step of these experiments. Despite the presence of those abundant non-physiological reads, our method seems to have accurately attributed genes to specific neuron types in most cases.

**Figure 5.**
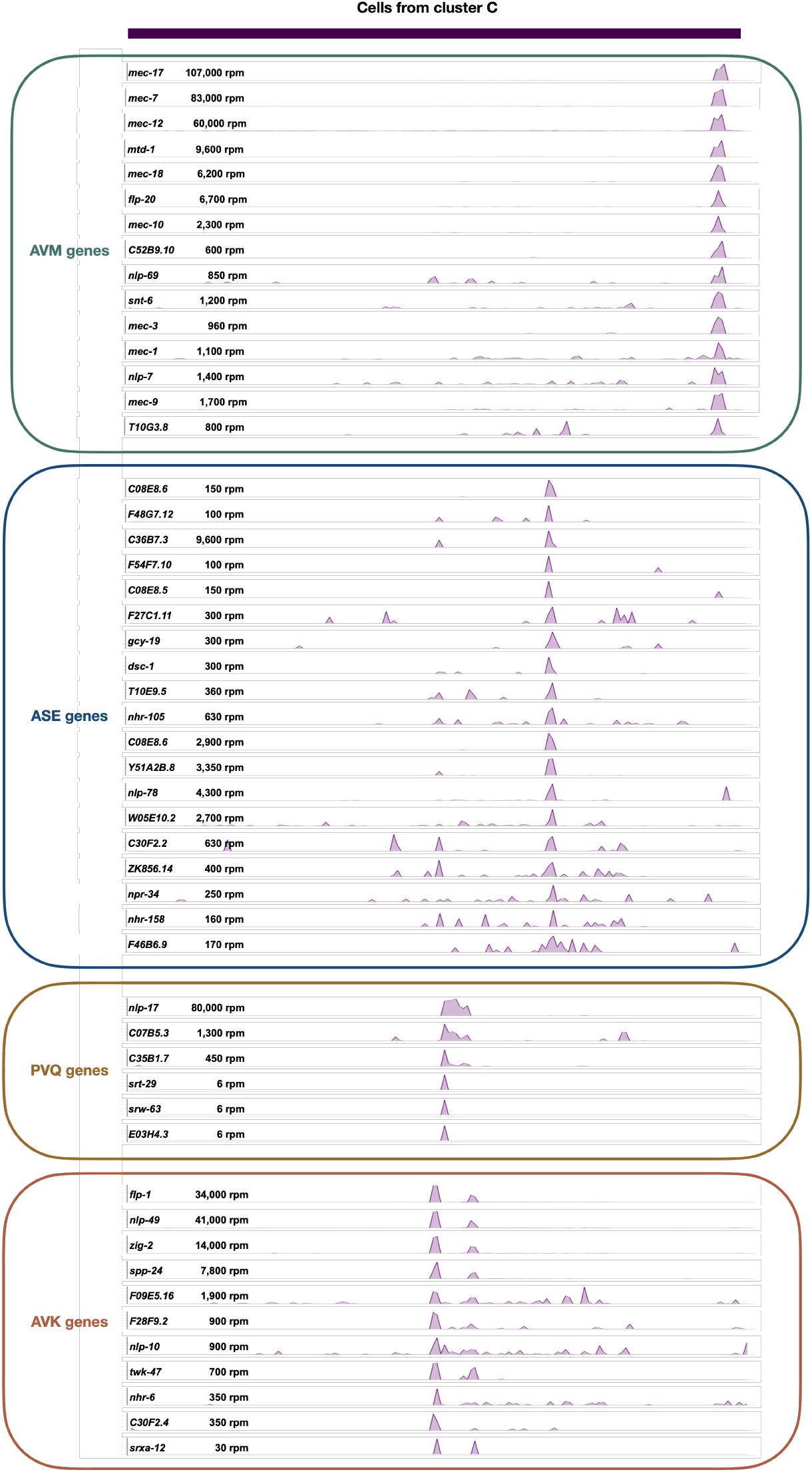
Read count plots for individual genes attributed to distinct neurons types (rpm) over the 164 High Coverage Cells from the “Neuronal” cluster C.

## DISCUSSION

The often overlooked limitation of current scRNA-seq technologies is that they only sequence a sample of the transcriptome of each cell^16^. We obtained accurate and detailed descriptions of molecular landscapes of individual cell types by using a subset of “High Coverage Cells” (HCC) contained in the tail end of published scRNA-seq data. By so reducing the dataset size, we could represent high content scRNA-seq using a classical gene vs cells expression heat map^24^. We devised a method to cluster genes that have strong specificity for cell groups identified by neighbor-joining.

Flared genes. The gene clusters we identified were highly specific and displayed a wide diversity of gene expression levels in their respective cells, including “flared” genes that can individually account for up to 10-20% of the reads in a given cell. In Excretory gland cells for example, ~27 genes account for >90% of detected transcripts; in the coelomic system, 90 genes account for ~50% of transcripts. Yet, other genes in their respective gene clusters show similar cell specificity but with expression levels sometimes a thousand fold lower. This observation raises fundamental questions on the mechanisms that allow genes to be expressed with similar cell specificity but with such different mRNA levels. In addition to the mechanistic question of how such levels of expression can be reached, the functionality of such accumulation will need to be investigated. For instance, why do embryonic intestinal cells need to have 10-20% of their mRNAs encoding TTR-50 (and another 10% of mRNAs from the 150 other genes in the *ttr-50* cluster) specifically during a 5 hour period during morphogenesis? The extent to which transcript levels correlate with the abundance of the corresponding protein is an often debated question. It will be very interesting to investigate if the flared genes identified here produce similarly high level proteins. If such amounts of proteins are indeed produced, it will provide new understanding of cellular functions and identities ; if they are not, it will raise questions regarding the function of the observed mRNA accumulations.

### Tissue Enrichment Ontology

We have obtained very significant tissue enriched annotation for clusters of cells that covers most of the animal tissue types and we could even resolve different states of the same cells during embryonic development (Figure 2). It is, however, important to bear in mind that current Gene Ontology terms rely on pre-existing annotations that are most frequently inferred based on co-expression with small sets of marker genes that have been experimentally confirmed for a given tissue, or cell. More often than not, genes are associated with multiple tissues due to contradicting evidence from various experimental sources^15^. For the 15 individual genes we tested in this study, we found that genes that were associated with a given tissue are indeed uniquely expressed in that tissue. Nevertheless, further experimental validation is warranted to explore to what extent the annotations provided here can help generate disambiguated lists of genes with tissue-specific expression. We believe the analysis presented here will be immediately useful to generate hypotheses regarding the molecular processes that are the basis of the establishment and maintenance of cell identity. To this end, we release a companion online tool that will facilitate these investigations for the *C. elegans* community (https://hcc-explorer.streamlit.app/Gene_Expression_Explorer). By exploring both cell expression spectrum visualization and individual gene expression plots, it will be more practical to evaluate the validity of the automated annotations we generated and determine which genes are better suited for experimental follow-up.

### Towards comprehensive single-cell transcriptomes

Importantly, we performed this analysis on “shotgun” datasets that were not specifically designed to reach saturation for the coverage of individual cells. We, however, only used <8% of the reads generated in these experiments. It is therefore likely that, despite using 15,000 rpm as a lower threshold of coverage (30 fold higher than the current standard), we are still only looking at a sampling of the cell mRNA content even for cells where hundreds of thousands of UMIs were collected. At this level of coverage we could detect genes with specific cellular levels of expression across 3 orders of magnitude (from hundreds of rpms to hundreds of thousands of rpms). Yet, it remains unclear if this covers the whole range of expression levels that could be found with a complete image of the mRNA landscape. To comprehensively characterize cellular transcriptomes, including the least expressed tier of cell-specific genes, it will therefore be necessary to adapt experimental strategies towards sequencing a lower number of cells in favor with a higher depth of transcriptome coverage.

## METHODS

### Raw data acquisition

Paired-end FASTQ files from publicly available *C. elegans* single-cell RNA-seq datasets were downloaded from the Sequence Read Archive (SRA) using the SRA Toolkit (v3.1.1). For each dataset, the study of origin and the different GEO accessions numbers associated with that study are listed in **Table 1**.

### Cell barcode and UMI extraction

Cellular barcode and UMI were extracted using an in-house python script based on fixed position accordingly with 10x Genomics Chromium Single Cell 3’ library type used for generating each dataset (v2 or v3).

### Reads alignment

cDNA-containing reads were aligned to *C. elegans* reference transcriptome, derived from the WBcel235/ce11 genome assembly and WormBase release WS295 (https://wormbase.org/) using Bowtie2 (v2.5.4, --fast-local mode). Alignment files were converted into BAM files using Samtools (v1.21).

### Generation of cell gene count matrix

For each dataset, the extracted cellular barcode, UMI, and gene alignment information were merged at the read level to generate a gene-UMI count table for each unique cellular barcode. Reads containing the same cellular barcode were then deduplicated by collapsing them into unique molecule counts per gene in order to produce a raw cell gene count matrix for each dataset. Cells with ≥15,000 total mapped UMIs (High Content Cells) were retained for the main analysis and merged into a single raw matrix.

### rpm and vmax normalization

The raw cell gene count matrix was normalized to reads per million (rpm) to account for sequencing depth by scaling each gene count to the total UMI count within a given cell, expressed on a per-million basis. For visualization purpose, a vmax-normalized matrix was generated from the rpm-normalized counts by dividing each gene count to its maximum rpm value across all cells.

### Unsupervised cell and gene clustering

Cells and genes were initially independently hierarchically clustered using Ward’s linkage method with Euclidean distance (SciPy v1.15.2).

### Supervised gene clustering

For each cluster of the cell hierarchical tree, we constructed a synthetic expression profile. This profile assigns a value of 100 to cells belonging to that node and 0 to all remaining cells. Each gene expression profile was then compared to each synthetic node expression profile and their similarity was assessed by computing a Pearson Correlation Coefficient score (PCC), resulting in a PCC scores matrix of ‘cell groups’ (nodes in the NJ-tree) versus genes. Each gene was then assigned to the node with which it correlated most strongly (highest PCC) in order to generate as many cluster-specific gene lists as possible. When applicable, cluster nodes were labeled by their most dominant gene, defined as the gene with the highest mean expression among the cells of the cluster.

### Tissue enrichment annotation

Cluster annotations were done by feeding cluster-specific gene lists into a local installation of the Wormbase Tissue Enrichment Analysis tool (v0.13.13)^15^ in order to identify cell types significantly over-represented within each list. We assigned the top hit as the cluster tissue annotation (p-adjusted value < 0.005). Clusters for which we could not identify a clear annotation remained unannotated.

### Collection and preparation of embryos for single-molecule fluorescent in situ hybridization

This step was performed according to the following protocol: https://dx.doi.org/10.17504/protocols.io.8epv52m74v1b/v1 ^16^. Wild-type (N2) worms were cultivated at 20°C on nematode growth medium (NGM) plates seeded with *E. coli* OP50 under standard laboratory conditions. Gravid adults were collected from the plates with M9 buffer and treated with hypochlorite solution. The embryos were washed in M9 buffer and filtered through a 40µm nylon filter. The embryos were fixed in fixation buffer (1X PBS, 37% Formaldehyde, H^2^O) at room temperature under rotation, washed in PBS solution containing 0.01% Tween-20, and stored in 70% ethanol at 4°C for four days. Prior to hybridization, embryos were permeabilized with 0.2 U/mL of chitinase for 5min at room temperature and washed with PBS solution containing 0.01% Tween-20.

### Single-molecule fluorescent in situ hybridization (smFISH)

smFISH was performed as described by Tsanov and collaborators^17^, using unlabelled primary probe pools ordered from Integrated DNA Technologies (IDT) and resuspended in pH 8.0 Tris-EDTA (TE) buffer at a final concentration of 0.833µM/probe and pre-hybridized with secondary probes Alexa Fluor 647 (AF647) labeled FLAP oligonucleotides. Permeabilized embryos were hybridized overnight at 37°C with the pre-hybridized probe-FLAP complexes, washed, and mounted in DAPI-containing medium prior to imaging.

### Image acquisition and processing

Images were acquired using a Zeiss Axio Imager Z.1 microscope equipped with a 100× objective. smFISH signal was acquired at 500ms exposure time and 20% laser intensity. DNA staining signal (DAPI) at 50ms exposure time and 20% laser intensity. Bright-field signal was acquired using 150ms exposure time. For each embryo, we acquired a Z-stack composed of 20 optical sections separated by 1 µm, for approximately five embryos per developmental stage and experimental condition. Maximum-intensity projections images from five consecutive optical sections were generated using ImageJ software.

## Supporting information

Video_abstract

## CODE AND DATA AVAILABILITY

The full implementation details of our analysis pipeline, including scripts and methods necessary to reproduce results and figures presented in this study will be made available via a code repository platform. Datasets tables and all other files generated as part of this study are available through an open-access repository platform (https://hcc-explorer.streamlit.app/Gene_Expression_Explorer).

## SUPPLEMENTAL MATERIAL

**Supplementary Figure 1:**
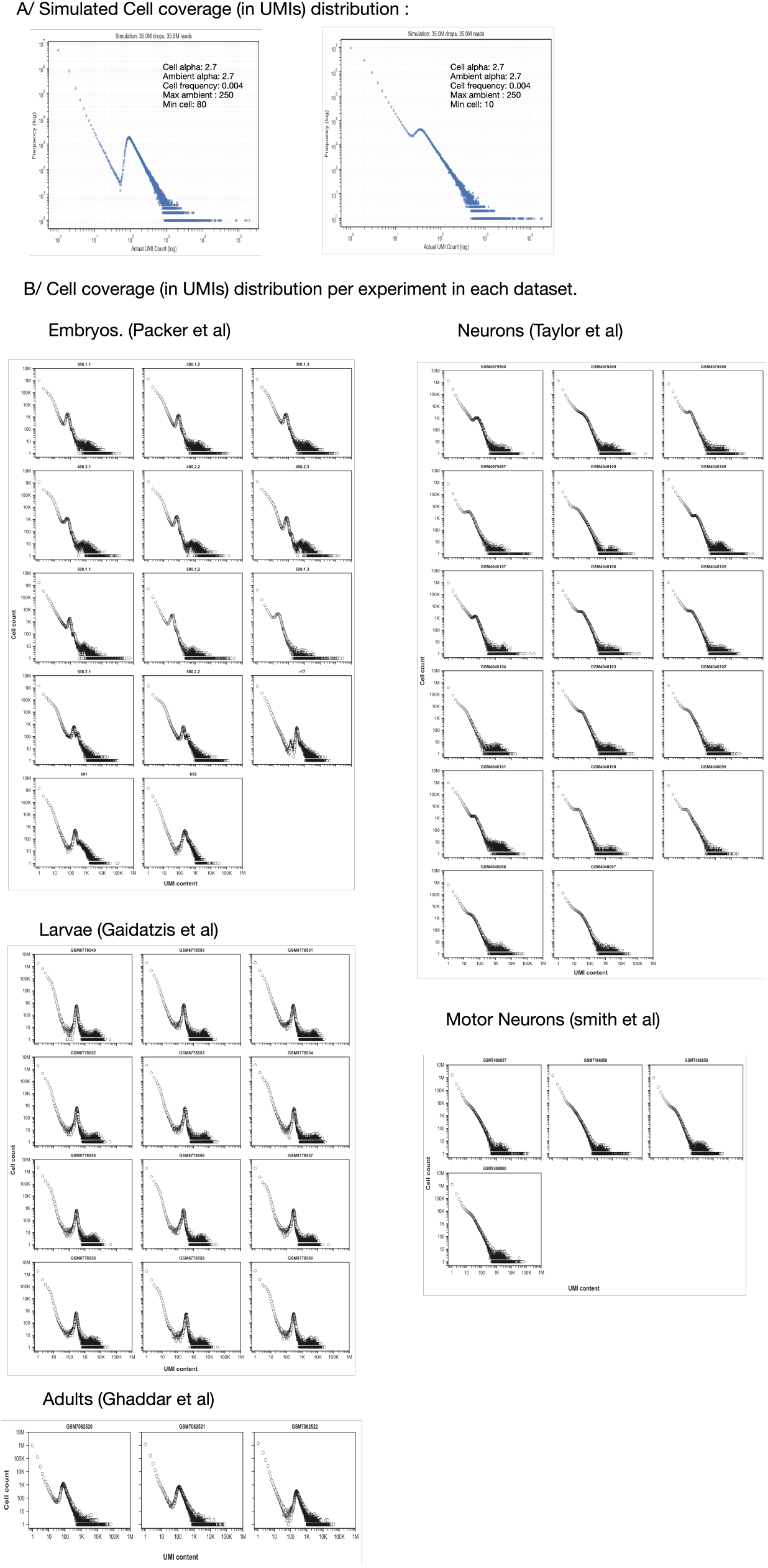
**A)** Simulated coverage distribution (see methods). **B)** UMI coverage distribution of individual sequencing experiments in the 5 published datasets used in this study.

## Notes

### Competing Interest Statement

The authors have declared no competing interest.

